# PhysioMotion Artifact: A Task-Driven EEG Dataset with Channel-Level Motion Artifact Annotations

**DOI:** 10.1101/2025.10.10.681755

**Authors:** Chunfeng Yang, Jiangwei Yu, Aonan He, Wentao Xiang, Xi Wang, Guangquan Zhou, Yudong Zhang, Miao Cao, Yang Chen, Juan M Gorriz

**Affiliations:** Key Laboratory of New Generation Artificial Intelligence Technology and Its Interdisciplinary Applications (Southeast University), Ministry of Education, Jiangsu Provincial Joint International Research Laboratory of Medical Information Processing, School of Computer Science and Engineering, Southeast University and Centre de Recherche en Information Biomédicale Sino-français (CRIBs), 2 Sipailou, Nanjing, 210096, Jiangsu, China; School of Biological Science and Medical Engineering, Southeast University, Nanjing, Jiangsu, 210096, China; Jiangsu Province Engineering Research Center for Smart Wearable and Rehabilitation Devices, School of Biomedical Engineering and Informatics, Nanjing Medical University, Nanjing, 211166, Jiangsu, China; School of Health Sciences, Swinburne University of Technology, John Street, Hawthorn, Victoria, 3122, Australia; Data Science and Computational Intelligence Institute, University of Granada, Granada, 52005, Spain

## Abstract

Physiological artifacts pose persistent challenges in electroencephalo-gram (EEG) data acquisition, often compromising interpretation and post-analysis of EEG signals across research and clinical applications. To address such limitations, including various artifact types, insufficient annotations, and low spatial resolutions, we present PhysioMotion Artifact, a large-scale, task-driven EEG dataset with channel-level artifact annotations. EEG data was acquired from 30 healthy participants performing 16 systematically designed single-type and multi-type movement tasks, inducing 14 distinct types of physiological artifacts. To demonstrate the utility of the dataset, we implemented a Convolutional Neural Networks-Transformer hybrid model for artifact detection and classification, achieving 98% accuracy in binary classification and 85% in 14-class classification tasks.

## Background & Summary

The EEG holds an irreplaceable position within the realms of neuroscience and neurological research. In addition to clinical diagnostics and assessing physiological states, EEG is crucial in evaluating consciousness levels like comas, and serves as an indispensable modality in the diagnosis and study of epilepsy1. However, acquiring EEG signals is frequently hindered by the presence of various physiological and non-physiological artifacts ^3^, including electrode displacement, movement artifacts, ocular artifacts, and artifacts originating from muscle activity. Such artifacts can significantly impede the accurate interpretation of EEG data^2^. Although proper environmental and equipment configurations can effectively reduce the level of non-physiological artifacts, physiological artifacts remain a major challenge in EEG data acquisition. Therefore, accurate and reliable detection and classification of EEG artifacts play a pivotal role in a wide range of applications, including clinical diagnosis and neuroscience research^4–8^, and brain-computer interface (BCI) systems^9–11^, where the reliability and interpretability of neural signals are critically dependent on signal quality.

In previous studies, artifact detection and classification have primarily relied on conventional signal processing techniques including Blind Source Separation (BSS) ^12–14^, Wavelet Transform^15–17^, and Empirical Mode Decomposition (EMD) ^18–20^. Those techniques offer advantages such as realtime applicability and the ability to extract artifact-related features with minimal computational cost. However, their effectiveness is often contingent upon expertise and manually optimized hyperparameters. In recent years, researchers have increasingly employed deep neural networks to automatically extract discriminative representations of various types of artifacts. Compared to traditional approaches, deep learning models, such as Convolutional Neural Networks (CNNs) ^21–23^, U-Net architectures^24;25^, and Transformer-based frameworks^26^, reduce dependence on handcrafted features and manual parameter tuning, while offering enhanced generalization across subjects and recordings. Despite these advantages, limited availability of high quality controlled and well-annotated EEG datasets remain as a major barrier.

Notably, three major multi-artifact datasets have been introduced: the Automatic Artifact Removal Dataset^27^, the EEG Artifact/Noise Dataset^28^, and the TUH EEG Artifact Corpus (TUAR) ^29^. The EEG dataset contaminated with artifacts^28^ includes multiple artifact types, however, it is limited to single-channel recordings, lacks a standardized experimental paradigm, and has a relatively small data volume, making it unsuitable for deep learning applications. In contrast, the dataset: Automatic Artifact Removal^27^ provides a sufficient amount of data for deep learning and features well-structured and standardized experimental designs. Nonetheless, its annotations are primarily timestamp-based, lacking the finer granularity of channel-specific annotations. The TUH EEG Artifact Corpus (TUAR) ^29^, built upon the large-scale clinical TUH EEG dataset^30^, offers extensive data with artifact annotations at the channel level and includes overlapped artifacts which enhances its applicability. However, as a clinical dataset, it presents several limitations, including label imbalance, insufficient diversity of artifact types, and a lack of targeted sample selection. Additionally, since the annotations are manually generated, labeling errors are difficult to avoid.

Addressing the critical need for a large-scale EEG dataset with highquality annotations, diverse artifact types, and high spatial resolutions, we designed a standardized experimental paradigm informed by the strengths and limitations of existing datasets. Task-driven EEG data were collected from 30 healthy participants, encompassing 14 distinct artifact types across two categories: (1) single-type tasks and (2) multi-type tasks. To synchronize task conduction with EEG recordings, we developed a custom data acquisition system that automatically logs task onset and offset. Signals were acquired using longitudinal and transverse bipolar montages standardized by the American Clinical Neurophysiology Society^31^. Artifact annotations, performed at the channel level, combined automated logging with manual validation to maximize annotation accuracy. To our knowledge, this represents the first publicly available large-scale EEG artifact dataset featuring task-based triggers and channel-specific annotations. The dataset is hosted on OpenNeuro (https://openneuro.org/crn/reviewer/eyJhbGciOiJIUzI1NiIsInR5cCI6IkpXVCJ9.eyJzdWIiOiIxYzI3MTg3Yi03OWYxLTRkODctYTQzMS1hZjhjNzkxNDUxMzUiLCJlbWFpbCI6InJldmlld2VyQG9wZW5uZXVyby5vcmciLCJwcm92aWRlciI6Ik9wZW5OZXVybyIsIm5hbWUiOiJBbm9ueW1vdXMgUmV2aWV3ZXIiLCJhZG1pbiI6ZmFsc2UsInNjb3BlcyI6WyJkYXRhc2V0OnJldmlld2VyIl0sImRhdGFzZXQiOiJkczAwNjM4NiIsImlhdCI6MTc1MDg5NjgwMCwiZXhwIjoxNzgyNDMyODAwfQ.c2a-3DRyC7SYof5NA1Gg5DH7Nr9lbq43h3lf-niZs30). Accompanying code for data is openly accessible on GitHub (https://github.com/JiangweiYu221/PhysioMotion_Artifact).

To evaluate the dataset, we implemented a deep learning model to perform artifact detection and classification tasks. The artifact detection task achieved an average accuracy of 98% and the artifact classification (14 classes) yielded an average accuracy of 86%. Our results suggest that the proposed dataset provides a solid foundation for deep learning-based artifact detection and classification. We anticipate that this dataset will address existing challenges in artifact identification, classification and removal in EEG signals, and facilitate the advancement of deep learning methods in this domain.

## Methods

### Participant Description

A total of 30 healthy individuals (18 males and 12 females; age range: 19–25 years; mean age: 21.5 ± 2.0 years) were recruited through campus advertisements at Southeast University, Nanjing, China. Prior to participation, all individuals were provided written informed consent in accordance with institutional guidelines. Screening procedures were used to exclude participants with history of any neurological disorders or mental health issues. Data acquisition was conducted between 7 November 2024 and 16 January 2025. Detailed demographic information and the timeline of data acquisition are presented in Table 1. The study received ethical approval from the Ethics Committee of Nanjing Medical University (No.2022784) and was conducted in strict accordance with the principles outlined in the Declaration of Helsinki.

**Table 1:** Information of 30 healthy participants (12 females and 18 males)

| Subject.ID | Gender | Age | First.Collection.Date | Second.Collection.Date |
| --- | --- | --- | --- | --- |
| 1 | male | 24 | Nov. 7, 2024 | Nov. 8, 2024 |
| 2 | male | 19 | Nov. 14, 2024 | Nov. 20, 2024 |
| 3 | male | 19 | Nov. 15, 2024 | Nov. 16, 2024 |
| 4 | male | 20 | Nov. 19, 2024 | Nov. 22, 2024 |
| 5 | male | 23 | Nov. 21, 2024 | Nov. 26, 2024 |
| 6 | male | 19 | Nov. 25, 2024 | Dec. 1, 2024 |
| 7 | male | 19 | Nov. 29, 2024 | Nov. 30, 2024 |
| 8 | male | 19 | Dec. 2, 2024 | Dec. 3, 2024 |
| 9 | male | 21 | Dec. 4, 2024 | Dec. 5, 2024 |
| 10 | female | 21 | Dec. 6, 2024 | Jan. 13, 2025 |
| 11 | female | 21 | Dec. 6, 2024 | Dec. 14, 2024 |
| 12 | male | 20 | Dec. 7, 2024 | Dec. 12, 2024 |
| 13 | female | 24 | Dec. 9, 2024 | Dec. 16, 2024 |
| 14 | female | 23 | Dec. 10, 2024 | Dec. 13, 2024 |
| 15 | female | 22 | Dec. 14, 2024 | Jan. 9, 2025 |
| 16 | female | 19 | Dec. 17, 2024 | Dec. 26, 2024 |
| 17 | female | 20 | Dec. 18, 2024 | Dec. 28, 2024 |
| 18 | female | 21 | Dec. 18, 2024 | Dec. 24, 2024 |
| 19 | female | 24 | Dec. 26, 2024 | Dec. 26, 2024 |
| 20 | female | 19 | Dec. 27, 2024 | Dec. 28, 2024 |
| 21 | female | 19 | Dec. 28, 2024 | Dec. 29, 2024 |
| 22 | male | 23 | Dec. 30, 2024 | Dec. 30, 2024 |
| 23 | male | 25 | Dec. 30, 2024 | Jan. 16, 2025 |
| 24 | male | 23 | Jan. 6, 2025 | Jan. 14, 2025 |
| 25 | male | 23 | Jan. 10, 2025 | Jan. 10, 2025 |
| 26 | male | 23 | Jan. 12, 2025 | Jan. 12, 2025 |
| 27 | female | 22 | Jan. 13, 2025 | Jan. 13, 2025 |
| 28 | male | 24 | Jan. 15, 2025 | Jan. 15, 2025 |
| 29 | male | 22 | Jan. 15, 2025 | Jan. 15, 2025 |
| 30 | male | 24 | Jan. 16, 2025 | Jan. 16, 2025 |

### Experimental Paradigm

The experimental protocol was designed to investigate individual and concurrent EEG artifacts through a structured sequence of movement tasks, which is motivated by the Temple University Artifact Corpus (TUAR) ^29^.

All tasks were conducted under the automated control of a custom-designed data acquisition system. This system utilized a combination of visual cues, textual prompts, and auditory instructions to provide real-time guidance, thereby facilitating precise execution of the pre-defined movements. Moreover, the system was programmed to automatically generate task-specific event markers and initiate countdown timers in alignment with pre-established temporal parameters.

The single-type tasks consisted of two baseline recordings, three oculomotor tasks, two head movement tasks, one tongue movement task, one chewing task, one swallowing task, and one eyebrow movement task. The experimental procedure was structured as follows: the session began with a 2-minute eyes-closed resting state, which served as a baseline with minimal artifacts. Subsequently, participants were prompted to blink their eyes freely and enter a 2-minute eyes-open resting state, which served as a second baseline recording. Following this baseline recording, oculomotor tasks were conducted. The first and second oculomotor tasks required participants to follow a red visual cue displayed on the screen, guiding vertical and horizontal eye movements, respectively. After some rest, the third oculomotor task required participants to blink continuously in every second for one minute. Head movement tasks were then carried out with participants’ eyes closed to avoid contamination by blink artifacts. Participants followed auditory cues to perform horizontal head rotations and vertical nodding. The tongue movement task also required eyes closed. Participants were instructed via auditory cues to repeatedly articulate the sound “la-la-la” ^32^ with a 2-second interval between each cue. In the masticatory task, participants closed their eyes and, following auditory prompts, clenched their molars to simulate chewing movements. For the swallowing task, participants closed their eyes and swallowed in response to auditory cues. Finally, the eyebrow movement task involved participants keeping their eyes closed and raising their eyebrows in response to an auditory cue, maintaining the raised position for 10 seconds.

The multi-type tasks consisted of five overlapping tasks designed to induce compound physiological artifacts. The first two tasks involved head movements performed with eyes open. These tasks followed the same procedures and timing as the two head movement tasks in the single-type condition, with the sole distinction being that participants were allowed to keep their eyes open. This modification was introduced to capture overlapping artifacts arising from simultaneous head movements and eye blinks. Following this, the blinking–eyebrow elevation task required participants to raise their eyebrows in response to an auditory cue and maintain the position, while simultaneously blinking once per second in response to separate auditory tones. This task was designed to induce overlapping artifacts from concurrent blinking and eyebrow movement. Next, the eyebrow elevation–tongue movement task mirrored the structure and timing of the tongue movement task in the single-type condition. Participants were instructed to keep their eyes closed throughout the task to avoid blink-related contamination. The key distinction in this task was that participants maintained sustained eyebrow elevation during each round of articulation, thereby generating overlapping artifacts from eyebrow and tongue movements. The final task—the eyebrow elevation–swallowing task—required participants to raise their eyebrows in response to an auditory cue and maintain the position for 5 seconds, during which they were instructed to swallow saliva once at any time within the interval.

The experimental paradigm was meticulously designed to elicit a comprehensive range of EEG artifacts through both single-type and multi-type movement tasks, enabling a systematic investigation into the individual and combined artifacts on EEG signals. All detailed information of experimental tasks is shown in Table 2.

**Table 2:** An overview of all tasks incorporated in the experimental paradigm, along with the corresponding manually annotated label names assigned to each task.

| Task Type | Task Description | Parameters | Notes | Label |
| --- | --- | --- | --- | --- |
| **Single-type Tasks** |  |  |  |  |
| Eyes-closed resting | Resting state | 2 mins duration | Baseline recording | close_base |
| Eyes-open resting | Resting state | 2 mins duration | Baseline recording with visual fixation | open_base |
| Vertical saccades | Ocular Control | 10 trials (5↑+5↓), 1 sec intervals | Instructed by the Red Dot | ver_eyem |
| Horizontal saccades | Ocular Control | 10 trials (5←+5→), 1 sec intervals | Instructed by the Red Dot | hor_eyem |
| Blinking | Metronomic blinking | 1 Hz frequency × 60s | Instructed by the Auditory cue | blink |
| Horizontal rotation (eyes closed) | Head Movement | 10 trials (5←+5→), 3 secs intervals | Instructed by the Auditory cue | hor_headm |
| Vertical nodding (eyes closed) | Head Movement | 10 trials (5↑+5↓), 3 secs intervals | Instructed by the Auditory cue | ver_headm |
| "Lalala" articulation | Lingual Movement | 3 blocks × 10 trials, 2 secs intervals | Instructed by the Auditory cue | tongue |
| Bilateral jaw clenching | Jaw Movement | 2 blocks × 5 trials, 2 secs intervals | Instructed by the Auditory cue | chew |
| Swallowing | Voluntary swallowing | 5 trials, 5 secs intervals | Instructed by the Auditory cue | swallow |
| Eyebrow elevation | Facial Control | 2 blocks × 10 secs duration | Instructed by the Auditory cue | eyebrow |
| **Multi-type Tasks** |  |  |  |  |
| Horizontal rotation (eyes open) | Combined Movement | 10 trials (5←+5→), 3 secs intervals | Visual fixation condition | blink_hor_headm |
| Vertical nodding (eyes open) | Combined Movement | 10 trials (5↑+5↓), 3 secs intervals | Visual fixation condition | blink_ver_headm |
| Sustained eyebrow elevation + blinking | Combined Movement | 2 blocks × 5 secs duration | Blinking by the Auditory cue | blink_eyebrow |
| Sustained eyebrow elevation + articulation | Combined Movement | 3 blocks × 10 trials, 2 secs intervals | | tongue_eyebrow |
| Eyebrow elevation + swallowing | Combined Movement | 2 trials × 5 secs duration | Single swallow for each trial | swallow_eyebrow |

### Data Acquisition

The experimental protocol was implemented under controlled laboratory environment to uphold methodological rigor and ensure the integrity of the collected data. All recording sessions took place in a quiet, dedicated environment, with only essential experimental apparatus present. Non-essential electronic devices were excluded from the setting to mitigate potential sources of electromagnetic noise.

As for visual stimuli, they were presented on a 23.8-inch IPS monitor (1920×1080 resolution, 16:9 aspect ratio) positioned at 60 cm viewing distance, resulting approximately 47.41° horizontal saccade angle and 27.75° vertical saccade angle of the human eye. Further hardware details are shown in Table 3.

**Table 3:**
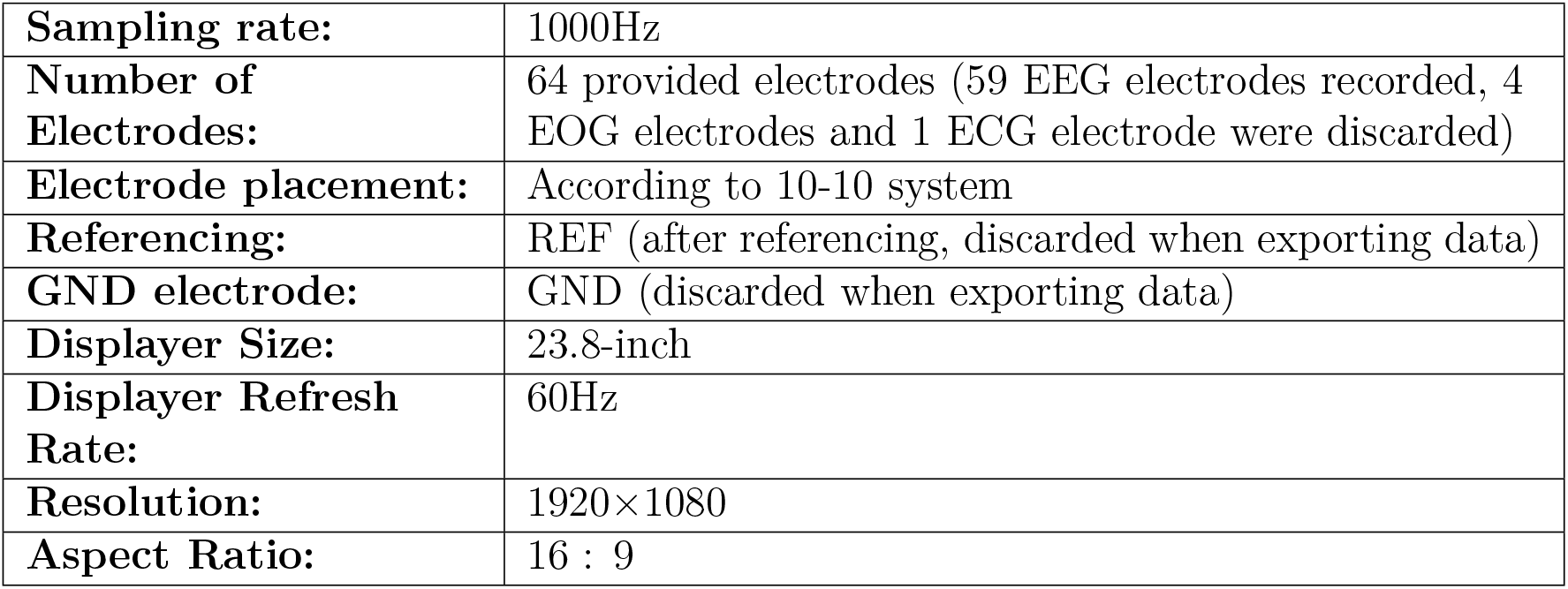
Hardware Specifications.

|  |  |
| --- | --- |
| **Sampling rate:** | 1000Hz |
| **Number of Electrodes:** | 64 provided electrodes (59 EEG electrodes recorded, 4 EOG electrodes and 1 ECG electrode were discarded) |
| **Electrode placement:** | According to 10-10 system |
| **Referencing:** | REF (after referencing, discarded when exporting data) |
| **GND electrode:** | GND (discarded when exporting data) |
| **Displayer Size:** | 23.8-inch |
| **Displayer Refresh Rate:** | 60Hz |
| **Resolution:** | 1920×1080 |
| **Aspect Ratio:** | 16 : 9 |

The visual cues consisted of red dot targets (20-pixel radius) containing 2-pixel-wide black crosshairs, optimized for precise oculomotor tracking. The complete set of four fixation cues used to elicit vertical and horizontal eye movements is illustrated in Figure 1.

**Figure 1:**
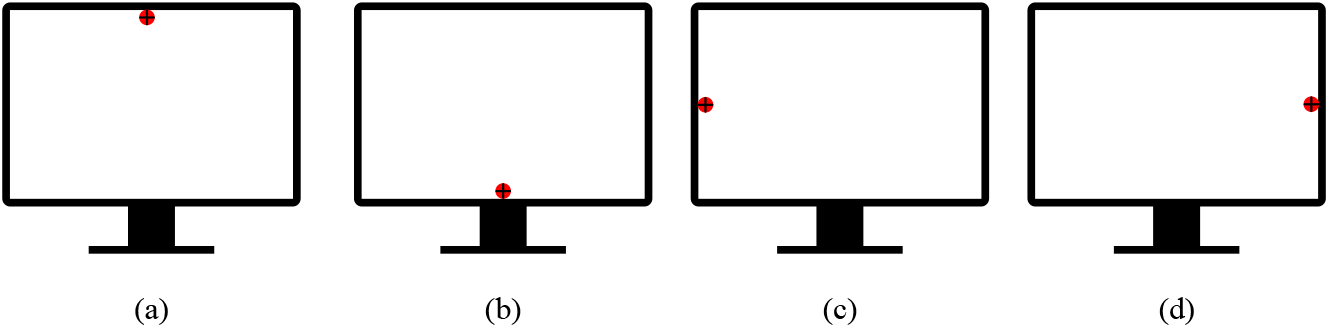
Visual Cues for Vertical Saccades (a&b) and Horizontal Saccades (c&d)

Once the experimental environment and equipment were configured as depicted in Figure 2, the formal data acquisition procedure was initiated. Within the designated acquisition room, each participant was seated comfortably in an upright posture facing the visual display monitor. To minimize potential electromagnetic interference from essential electronic components, the acquisition laptop and wireless router were positioned at a distance of approximately 1.5 meters from both the participant and the prompt display system.

**Figure 2:**
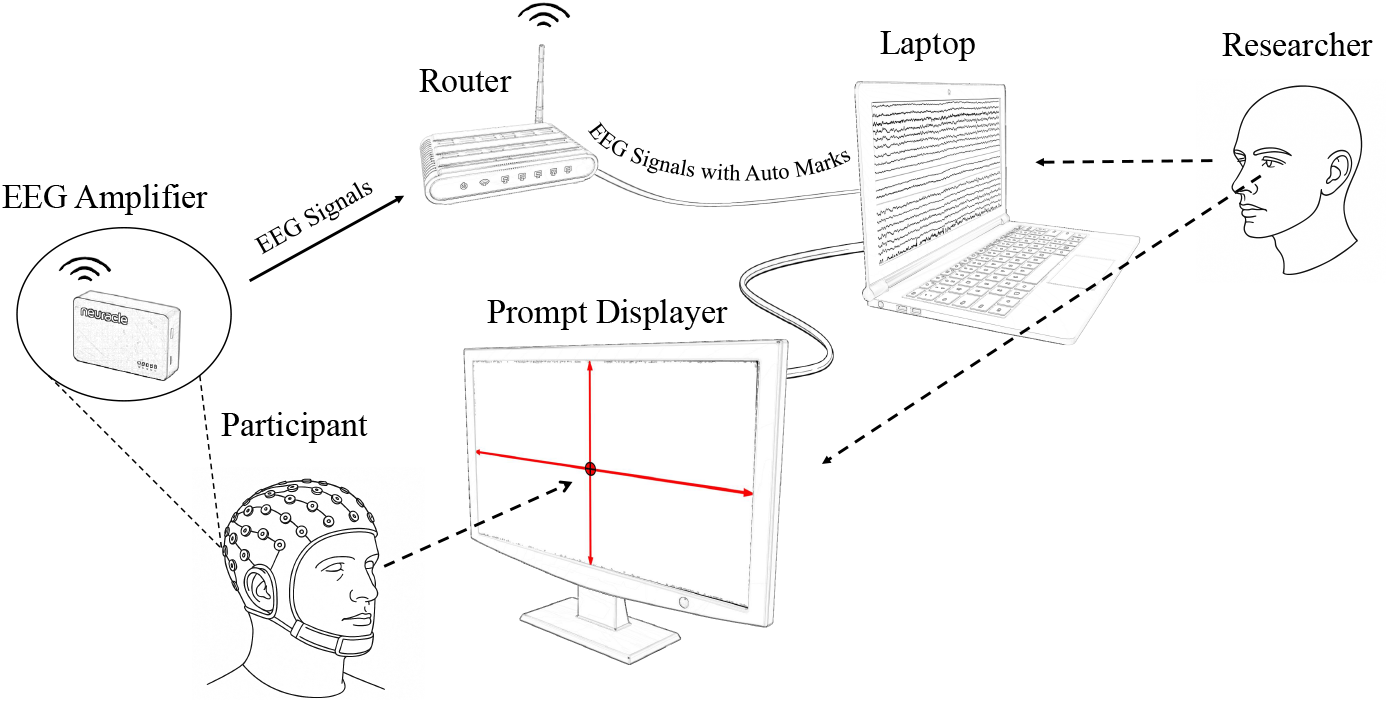
Data Acquisition Systems

Throughout recordings, the acquisition system provided real-time visualizations of the EEG signals, allowing the researcher to monitor data quality and verify the temporal alignment between task execution and recorded brain activity. Upon completion of the experimental session, raw EEG data were exported from the acquisition laptop and securely stored for subsequent analysis.

To accommodate individual differences in physical endurance and personal scheduling constraints, the experimental protocol incorporated a degree of flexibility. Participants were given the option to complete all six blocks either within a single day or over two days. For the two-day option, participants performed three blocks per day, with the specific dates determined based on their availability and preference. This flexible scheduling framework was designed to minimize discomfort, sustain high-quality data acquisition, and uphold ethical standards by prioritizing participants’ well-being. As a result of this design, all participants successfully completed the full set of experimental procedures, contributing a consistent dataset comprising approximately 120 minutes of EEG recordings per individual.

### Preprocessing and Annotations

In addition to the raw EEG recordings, the dataset also provides a corresponding set of preprocessed data. Given the central objective of this study—namely, the characterization of EEG artifacts—the preprocessing pipeline was designed to be minimal. A notch filter (50 Hz and its harmonics up to 150 Hz) was first applied to the raw EEG signals. Subsequently, a bandpass filter between 0.5 Hz and 150 Hz was adopted. This filtering strategy was designed to retain the temporal and spectral features of artifact-related components. Following filtering, both standard longitudinal and transverse bipolar montages were applied to EEG signals, in accordance with the recommendations of the American Clinical Neurophysiology Society^31^. Bipolar referencing was applied, enhancing the visibility of spatially localized artifact patterns. Details of Montages used in this study were shown in Table 4.

**Table 4:** Standard Longitudinal and Transverse Bipolar Montages (18.1).

| Channel No. | Standard Longitudinal Bipolar Montages-18.1 | Standard Transverse Bipolar Montages-18.1 |
| --- | --- | --- |
| 1 | Fp1-F7 | F7-Fp1 |
| 2 | F7-T7 | Fp1-Fp2 |
| 3 | T7-P7 | Fp2-F8 |
| 4 | P7-O1 | F7-F3 |
| 5 | Fp1-F3 | F3-Fz |
| 6 | F3-C3 | Fz-F4 |
| 7 | C3-P3 | F4-F8 |
| 8 | P3-O1 | T7-C3 |
| 9 | Fz-Cz | C3-Cz |
| 10 | Cz-Pz | Cz-C4 |
| 11 | Fp2-F4 | C4-T8 |
| 12 | F4-C4 | P7-P3 |
| 13 | C4-P4 | P3-Pz |
| 14 | P4-O2 | Pz-P4 |
| 15 | Fp2-F8 | P4-P8 |
| 16 | F8-T8 | P7-O1 |
| 17 | T8-P8 | O1-O2 |
| 18 | P8-O2 | O2-P8 |

Building upon preprocessed EEG signals, the subsequent artifact annotations were conducted in accordance with a rigorous quality control framework. Two experienced annotators independently performed channel-level artifact annotation under double-blind conditions to minimize subjective bias and enhance annotation reliability. Leveraging a customized labeling interface, and guided by system-generated auto markers, each annotator systematically identified the presence of target artifacts by selecting the corresponding EEG channels and signal segments. An overview of the preprocessing and annotation pipeline is presented in Figure 3.

**Figure 3:**
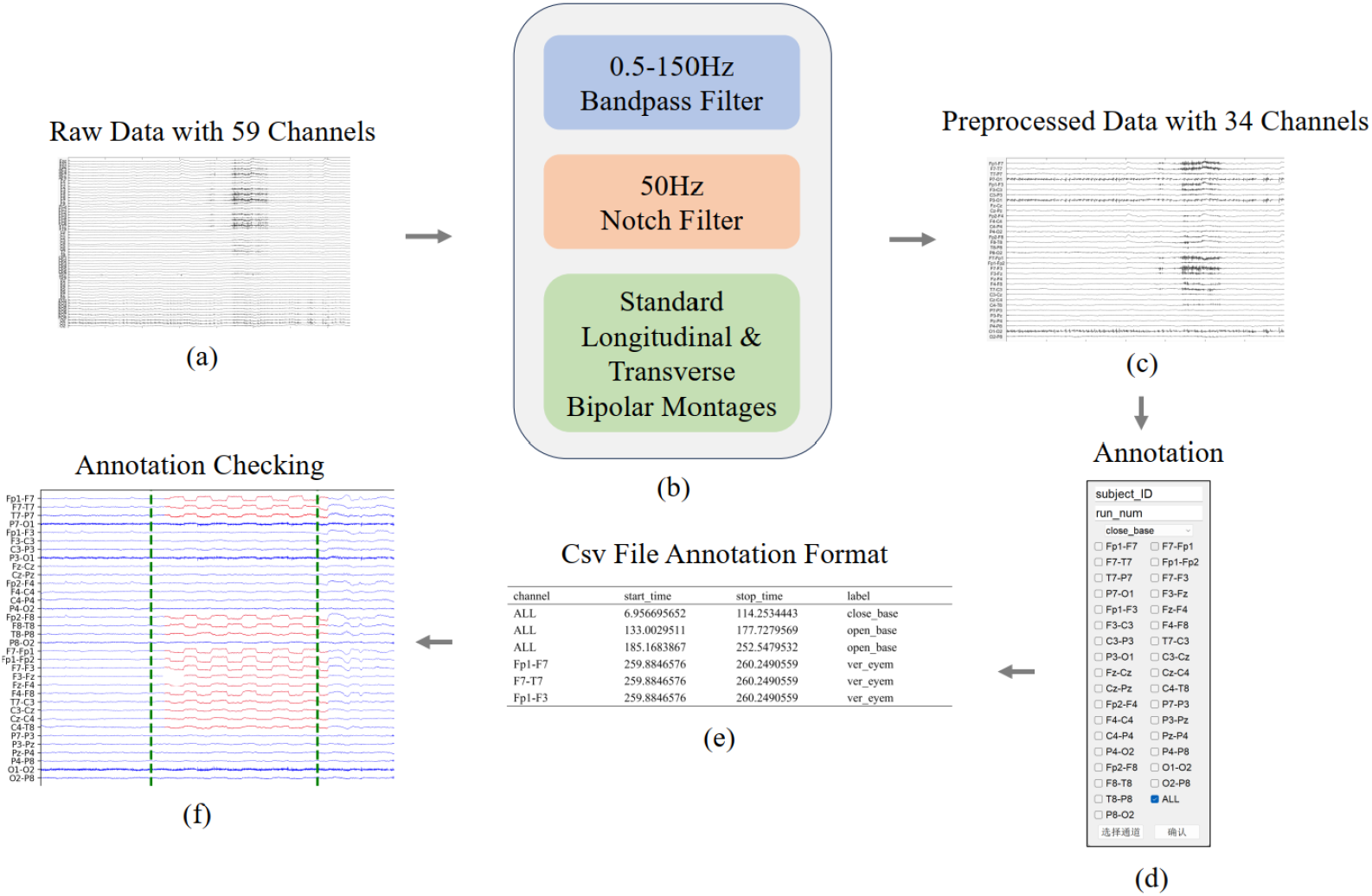
EEG Signal Processing and Annotation Pipeline. (a) Raw data with all 59 EEG channels exported from NeuSen W; (b) Three-step preprocessing including filtering and bipolar montage; (c) Preprocessed data with 34 montaged channels; (d) Customised annotation graphical user interface used for independent labeling; (e) Annotation data (f) Annotation verification interface for joint discussion.

## Data Records

The dataset is organized in full compliance with the Brain Imaging Data Structure (BIDS) standard33. Additionally, manually annotated artifact annotations are provided in csv files, which are stored in a dedicated folder named `Manual_Annotations`. The overall file organization and directory structure of the dataset are illustrated in Figure 4.

**Figure 4:**
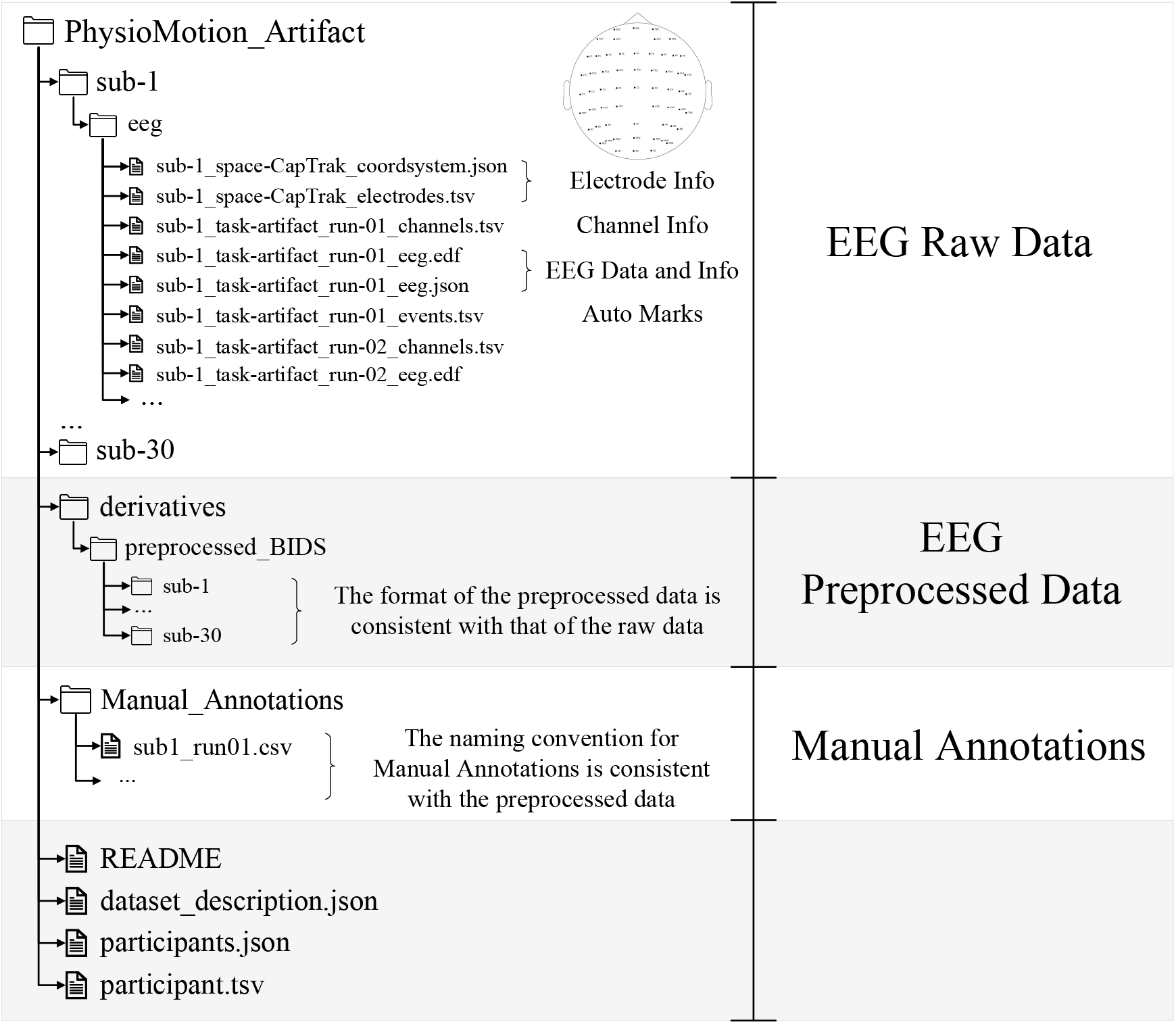
Directory and Details for the Dataset

### Raw Data

The raw EEG data in this study was recorded and exported using the NeuSen W system in.`bdf` files, with the signal units in microvolts (*µ*V). However, this format and unit convention pose limited compatibility with the MNE-Python^34^ framework, which operates with data in volts by default and is optimized for the BIDS format. To ensure consistency across the dataset, the signal units were converted from microvolts to volts, and the data were reorganized in full compliance with the BIDS specification. The resulting raw dataset strictly adheres to the BIDS format, following these conventions: the `subject` identifier ranges from `1` to `30`, corresponding to each participant; the `task` field is consistently labeled as `artifact`, reflecting the artifact-induction nature of the recordings; the `run` index ranges from `01` to `06`, indicating the number of experimental task rounds performed by each subject; and the `datatype` is specified as `eeg`. Each raw recording contains 59 EEG channels sampled at 1000 Hz.

### Preprocessed Data

All preprocessing procedures were implemented using the preprocessing functions from the MNE-Python^34^. Following the preprocessing pipeline, the resulting preprocessed data were saved in full compliance with the BIDS format, ensuring consistency with the raw data organization. The preprocessed EEG recordings contain 34 channels(after re-referencing) and retain the original sampling rate of 1000 Hz.

### Manual Annotations

The naming convention of the manually annotated data is kept consistent with that of the preprocessed data. Each annotation file follows the format:

`sub-x run0y.csv`

where, `x` denotes the subject ID ranging from 1 to 30, and y indicates the task session number from 1 to 6. Each CSV file contains four columns with the standardized headers: `channel, start_time, stop_time`, and `label`. A comprehensive summary of all channel-level manual annotation statistics is provided in Table 5.

**Table 5:**
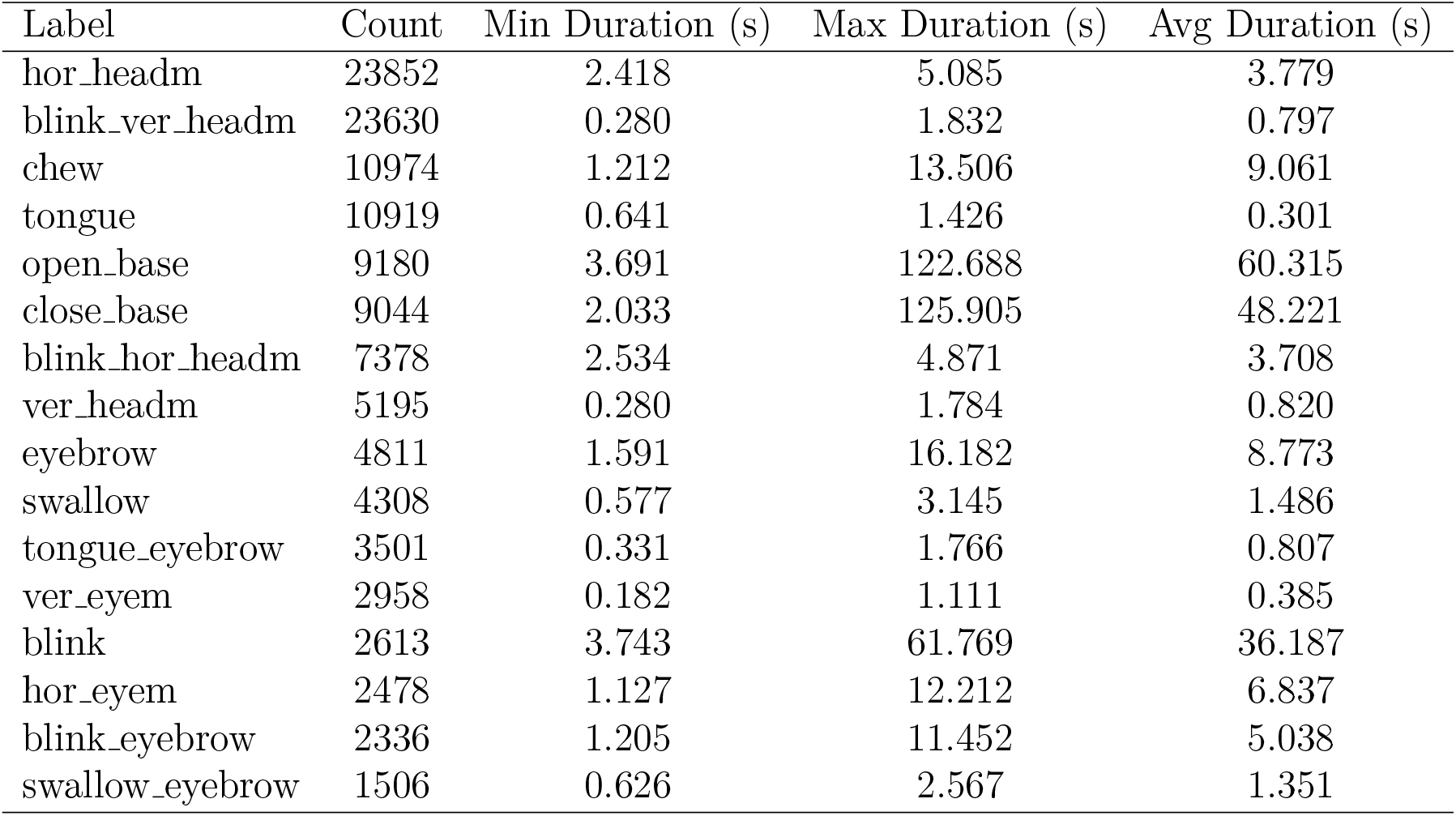
Statistical Information of All Manual Annotations (s represents the unit: second)

### Technical Validation

#### Topographic Mapping and Artifact Feature

As depicted in Figure 5, four representative EEG topographic maps alongside their corresponding signals were used to illustrate the distinct characteristics associated with different artifact types. These examples highlight the unique spatial distributions and temporal dynamics that generate each artifact category. Such patterns constitute critical features that deep learning models must capture in order to achieve robust artifact detection and classification.

**Figure 5:**
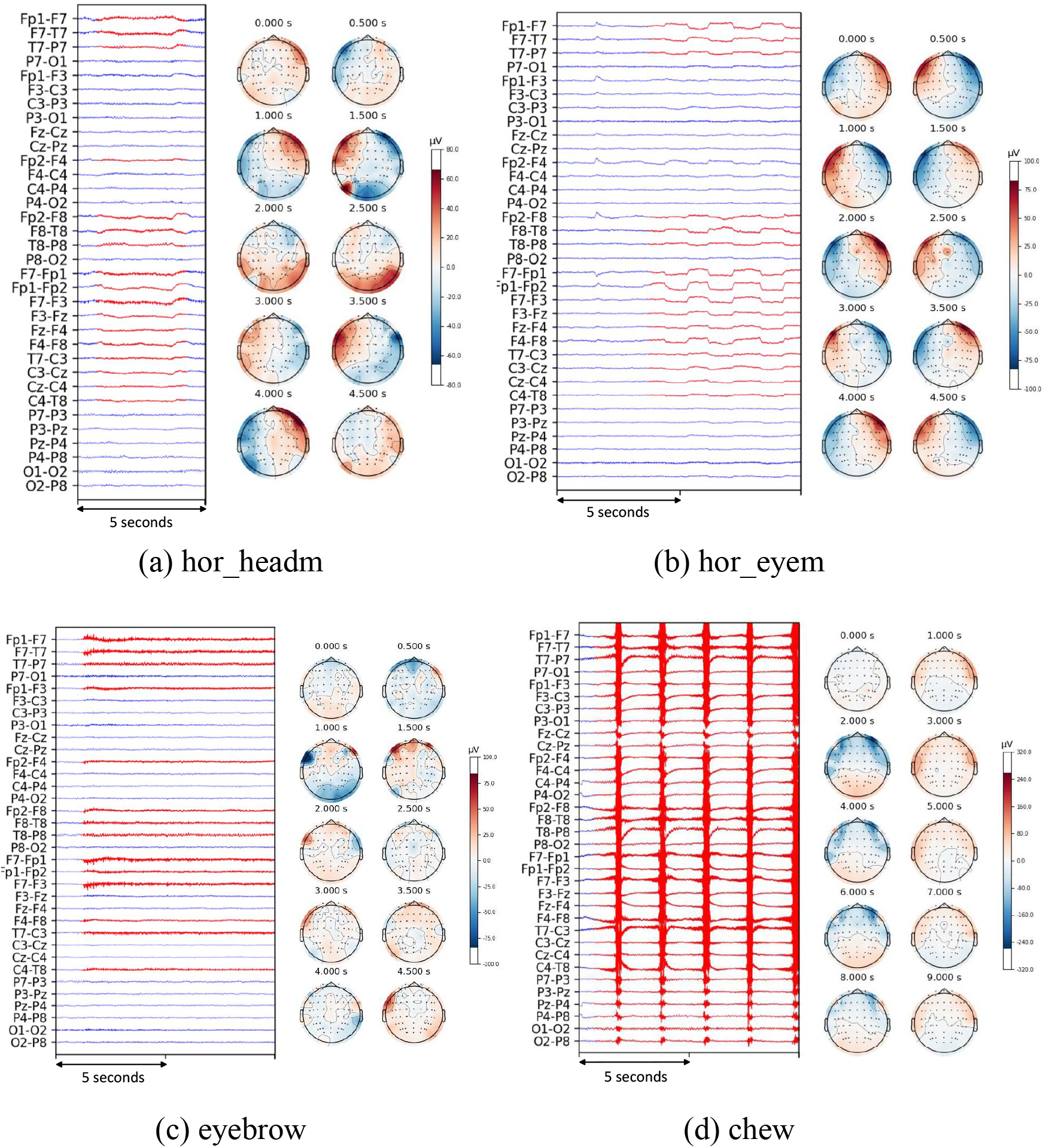
Directory and Details for the Dataset

Building upon this observation, the subsequent sections provide a comprehensive description of our artifact identification and classification framework, which is grounded in the use of deep neural networks.

#### Model Input

To improve computational efficiency and ensure integration with deep learning frameworks, preprocessed EEG signals were downsampled and segmented. Specifically, EEG signals were first downsampled to 125 Hz, a rate chosen to substantially reduce data dimensionality while retaining critical temporal features relevant for artifacts. After downsampling signals, continuous recordings were segmented into fixed-length 3-second time windows, corresponding to 375 time points per segment. To better capture temporal transitions across adjacent segments, a 50% overlap was implemented, leading to a sliding window stride of 187 samples. Moreover, to better utilise the training dataset and mitigate the inclusion of irrelevant or noise-dominated intervals, a targeted window selection strategy was adopted. Specifically, for each annotated artifact segment, the initial window was always retained, after which the sliding process proceeded in half-window increments until the segment extended beyond the labeled artifact boundary. This selective segmentation approach ensures that the model is predominantly exposed to windows enriched with clinically or analytically significant artifact patterns. Consequently, it enhances the model capacity to learn salient discriminative features and generalize effectively across varying artifact manifestations.

#### Model Architectures

To evaluate the dataset, we employed two structurally homologous CNN–Transformer classifiers inspired by Peh et al. ^26^ The binary classification (*K* = 2) detects artifact-contaminated EEG signals against resting-state signals, while the multiclass model (*K* = 14) classifies specific physiological artifact subtypes. Both models share a unified three-stage architecture: spatial feature extraction via 2D CNN, temporal modeling using Transformer encoders, and final classification through fully connected layers (FC). Given an input EEG data:

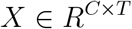

where *C* denotes the number of channels and *T* denotes the number of sample points. The prediction of the model can be expressed as,

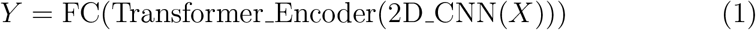

The overall CNN–Transformer architecture is illustrated in Figure 6.

**Figure 6:**
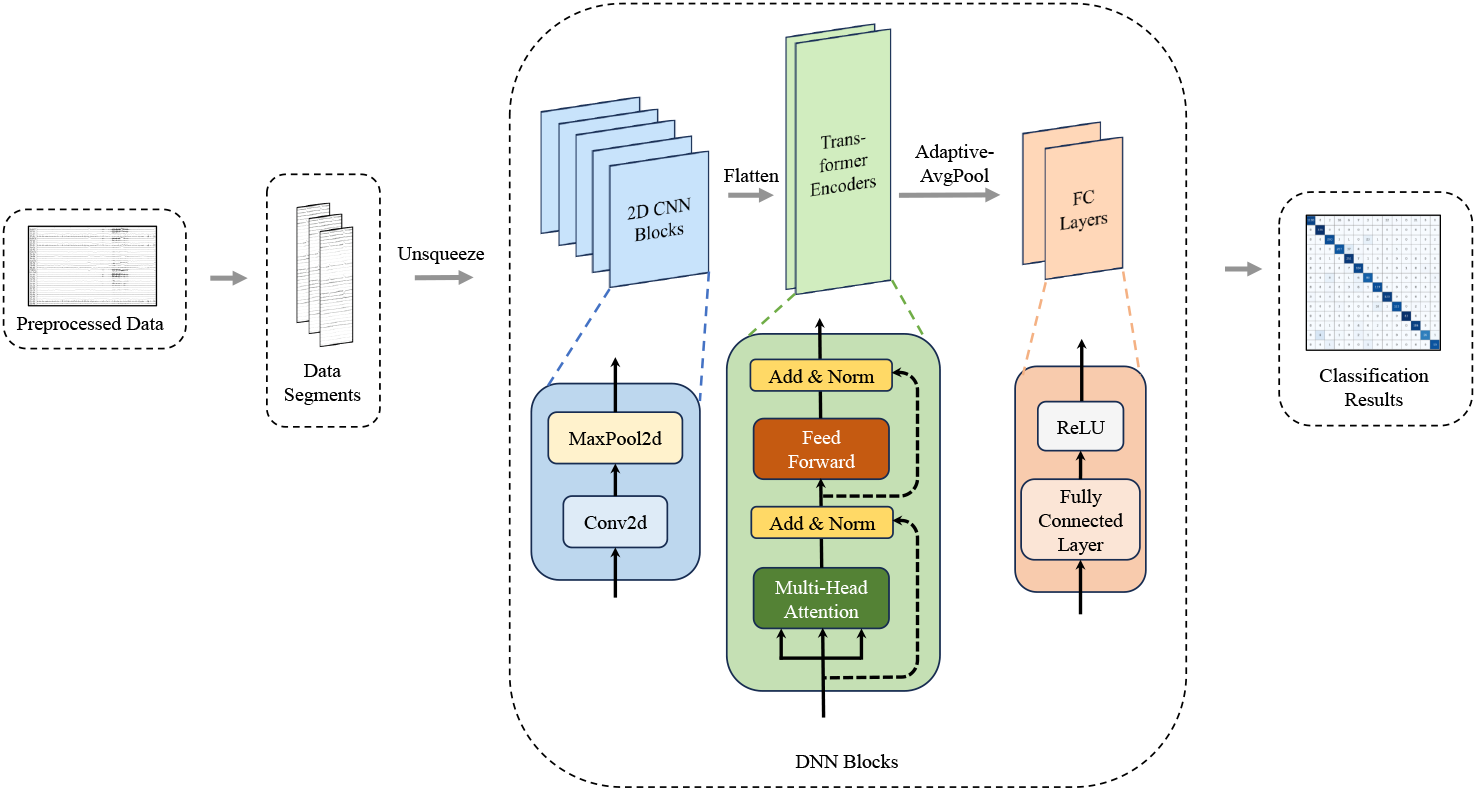
DNN Model Structure for Multi-Class Classification

Model parameters *θ* were optimized by minimizing a regularized class-weighted cross-entropy loss, which simultaneously addresses overfitting and class imbalance. The loss function of the model is defined as:

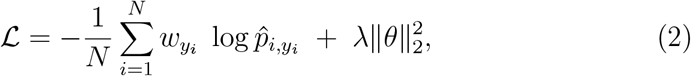

where *N* is the minibatch size, *y*_*i*_ *∈ {*1, …, *K}* denotes the ground-truth annotation, 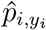 is the predicted probability, *w*_*yi*_ is the class-specific weight, and *λ* is the weight decay coefficient for L2 regularization. To mitigate the adverse effects of class imbalance, we adopted a two-stage strategy for computing class weights *w*_*k*_ used in the loss function. Initially, base weights 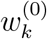 were derived from inverse class frequencies to proportionally upweight underrepresented classes:

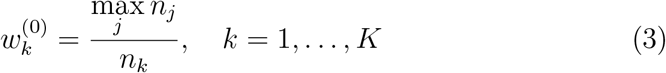

where *n*_*k*_ represents the number of training samples belonging to class *k*, and max_*j*_ *n*_*j*_ ensures that the most frequent class is assigned a baseline weight of 1. Subsequently, to refine this coarse adjustment and avoid overcompensation, we introduced adaptive scaling factors *s*_*k*_, yielding the final class weights:

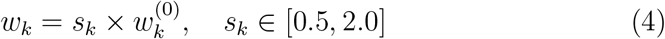

where *s*_*k*_ is a class-specific multiplier optimized via Hyperopt. The bounded range for *s*_*k*_ constrains the extent of adjustment, promoting stability during training. The resulting weights were integrated into the class-weighted crossentropy loss to guide model learning under imbalanced conditions.

#### Hyperparameter Optimization

We employed Hyperopt’s Tree-structured Parzen Estimator (TPE)35 to jointly tune learning rate, depth of the CNN, number of Transformer layers and attention heads, batch size, weight decay, and dynamic class-weight scalings. Table 6 summarizes the best configurations for each task.

**Table 6:** Best hyperparameters for binary and multi-class classifiers.

| Hyperparameter | Binary Task | Multi-Class Task |
| --- | --- | --- |
| Learning rate $\eta$ | 0.0009815 | 0.0003771 |
| Batch size | 128 | 128 |
| CNN blocks | 5 | 5 |
| Transformer layers | 1 | 2 |
| Attention heads | 16 | 16 |
| Weight decay | $4.35 \times 10^{-5}$ | $2.11 \times 10^{-6}$ |
| **Class-weight scaling $w_k$** |  |  |
| `close_base` | 1.78 | — |
| `artifact` | 1.84 | — |
| ... (14 entries) | — | see footnote * |

\* Multi-class weights (ordered by `blink`, `chew`, ..., `swallow_eyebrow`): {1.36, 1.26, 0.87, 1.44, 0.66, 0.79, 0.85, 1.03, 2.00, 1.00, 1.64, 0.69, 1.07, 1.82}.

## Results

The classification reports(Table 7) and confusion matrices(Figure 7) for both binary and multi-class tasks demonstrate that the CNN–Transformer models exhibit exceptional performance in artifact recognition on the proposed dataset. In the binary classification task, the model achieved a peak validation accuracy of approximately 99%, effectively distinguishing artifact-contaminated segments from baseline (artifact-free) segments with high precision. In the more challenging multi-class setting, the model attained an overall classification accuracy of approximately 92%, with a capacity of differentiating various artifact types. These results collectively highlight the robustness and generalizability of the CNN–Transformer architecture in performing both binary and multi-class artifact detection.

**Table 7:**
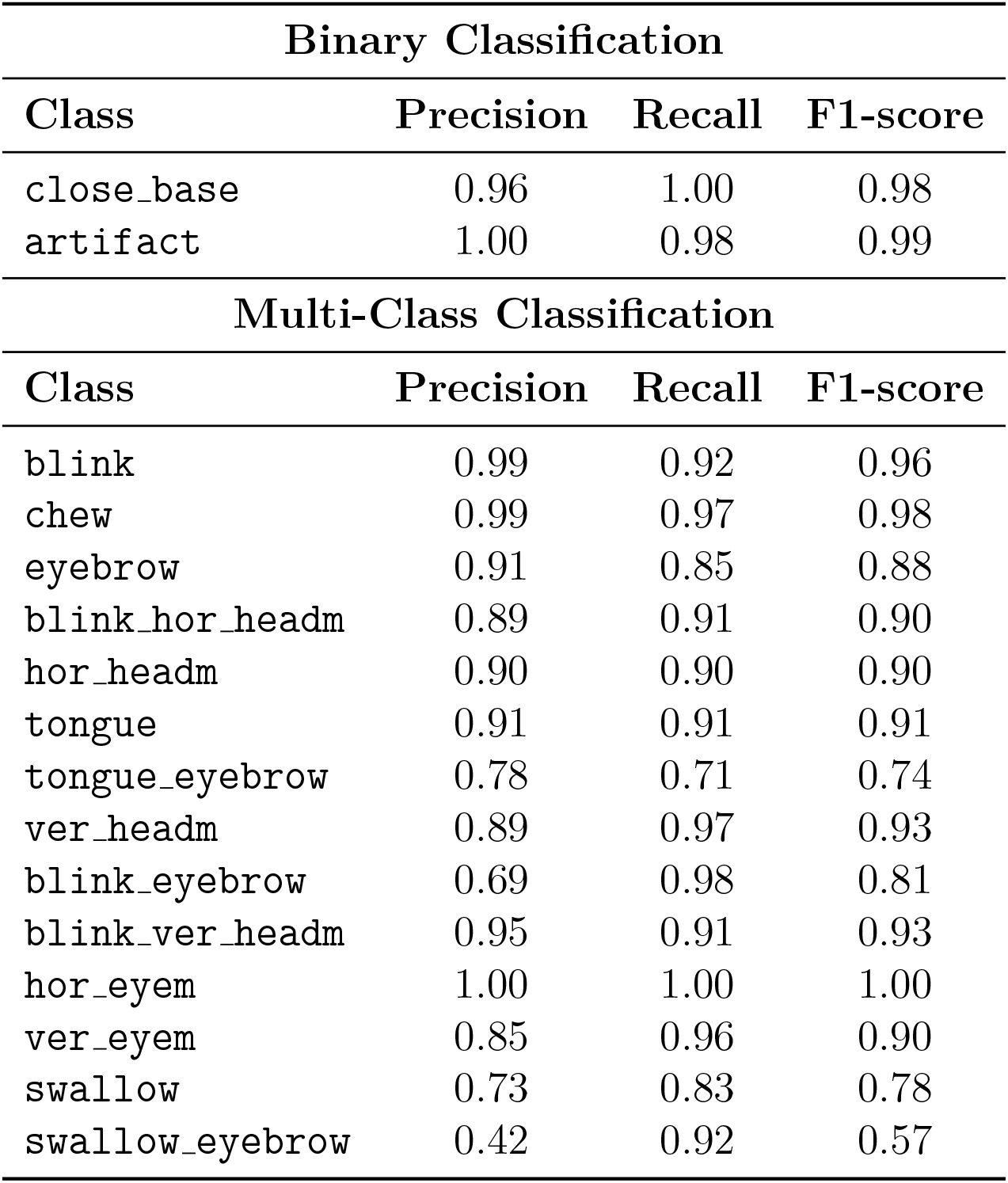
Precision, Recall and F1-score for Binary and Multi-Class Classification.

| Binary Classification |  |  |  |
| --- | --- | --- | --- |
| Class | Precision | Recall | F1-score |
| close_base | 0.96 | 1.00 | 0.98 |
| artifact | 1.00 | 0.98 | 0.99 |
| Multi-Class Classification |  |  |  |
| Class | Precision | Recall | F1-score |
| blink | 0.99 | 0.92 | 0.96 |
| chew | 0.99 | 0.97 | 0.98 |
| eyebrow | 0.91 | 0.85 | 0.88 |
| blink_hor_headm | 0.89 | 0.91 | 0.90 |
| hor_headm | 0.90 | 0.90 | 0.90 |
| tongue | 0.91 | 0.91 | 0.91 |
| tongue_eyebrow | 0.78 | 0.71 | 0.74 |
| ver_headm | 0.89 | 0.97 | 0.93 |
| blink_eyebrow | 0.69 | 0.98 | 0.81 |
| blink_ver_headm | 0.95 | 0.91 | 0.93 |
| hor_eyem | 1.00 | 1.00 | 1.00 |
| ver_eyem | 0.85 | 0.96 | 0.90 |
| swallow | 0.73 | 0.83 | 0.78 |
| swallow_eyebrow | 0.42 | 0.92 | 0.57 |

**Figure 7:**
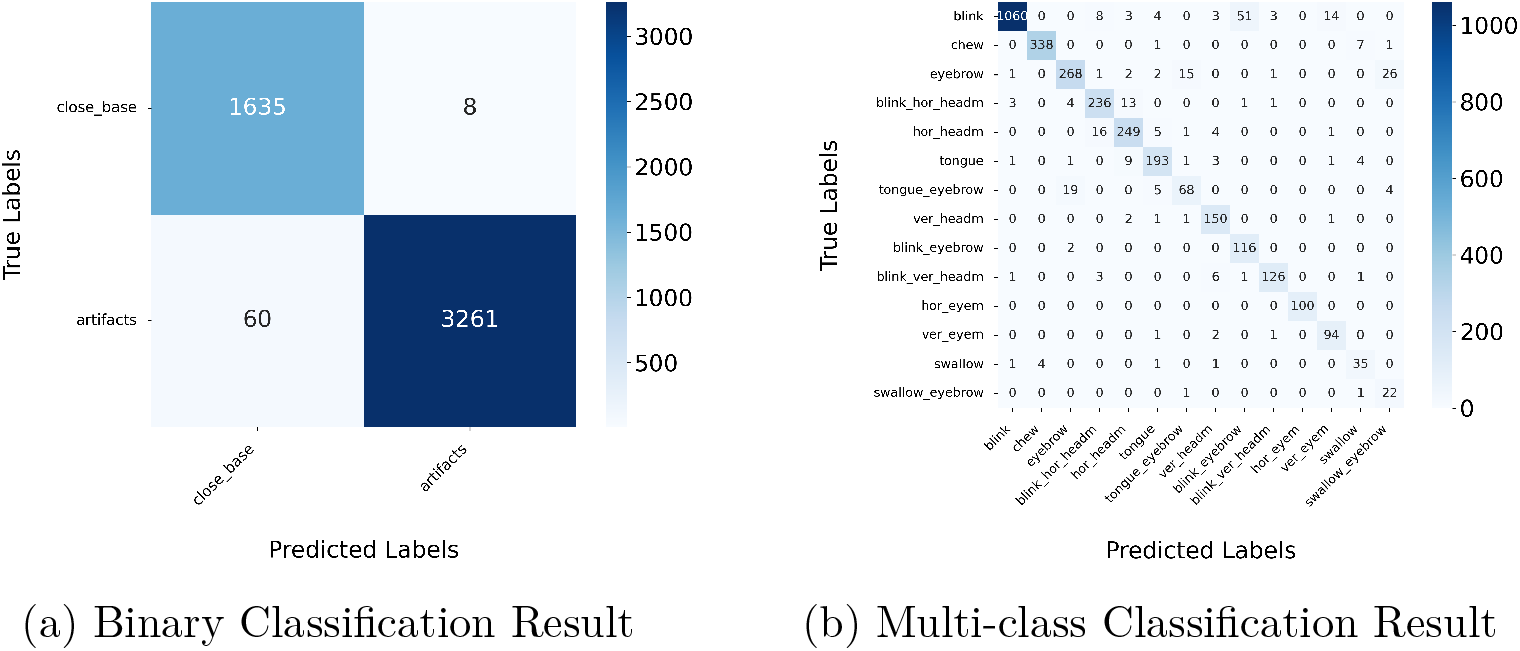
Confusion Matrix

To further quantify model performance, standard classification metrics—Precision, Recall, and F1-score—were calculated from the confusion matrices and are summarized in Table 7 for both binary and multi-class scenarios.

## Usage Notes

To facilitate the analysis and processing of EEG data within this dataset, Python 3.7 and the MNE-Python v1.3.0 were used. In the data acquisition system used for this study, automatic event marking was implemented via a proprietary software API36 provided by Neuracle Inc.

For the EEG artifact classification task, a hybrid model combining 2D Convolutional Neural Networks (CNNs) and Transformer layers was implemented using PyTorch. All experiments were conducted within a conda-managed environment running Python 3.8.20 on an Ubuntu-based Linux system equipped with two NVIDIA GeForce RTX 4090 GPUs (24GB each), using NVIDIA-SMI v565.57.01 and CUDA 12.7. The environment included the following key Python packages: PyTorch 2.4.1, NumPy 1.24.3, Scikitlearn 1.3.0, Hyperopt 0.2.7, GPUtil 1.4.0, TQDM 4.66.5, and the standard Python logging module.

## Code Availability

The majority of the code utilized in this study has been made publicly available on GitHub (https://github.com/JiangweiYu221/PhysioMotion_Artifact), encompassing modules for data preprocessing, the annotation interface, the annotation verification tool, and model training procedures. The data acquisition system mentioned in this work is not hosted in the repository but is freely available upon request by contacting the authors via email. Comprehensive instructions for setup and usage are provided in the accompanying README file included in the repository.

## Data Availability

The dataset is publicly available on OpenNeuro platform (https://openneuro.org/crn/reviewer/eyJhbGciOiJIUzI1NiIsInR5cCI6IkpXVCJ9.eyJzdWIiOiIxYzI3MTg3Yi03OWYxLTRkODctYTQzMS1hZjhjNzkxNDUxMzUiLCJlbWFpbCI6InJldmlld2VyQG9wZW5uZXVyby5vcmciLCJwcm92aWRlciI6Ik9wZW5OZXVybyIsIm5hbWUiOiJBbm9ueW1vdXMgUmV2aWV3ZXIiLCJhZG1pbiI6ZmFsc2UsInNjb3BlcyI6WyJkYXRhc2V0OnJldmlld2VyIl0sImRhdGFzZXQiOiJkczAwNjM4NiIsImlhdCI6MTc1MDg5NjgwMCwiZXhwIjoxNzgyNDMyODAwfQ.c2a-3DRyC7SYof5NA1Gg5DH7Nr9lbq43h3lf-niZs30).

## Acknowledgements

This work was supported by the National Natural Science Foundation of China under Grant T2225025. We also gratefully acknowledge the support provided by the Big Data Computing Center of Southeast University and Visiting Scholar programme at the UGR (Spain).

## Author Contributions

C.Y., G.Z., M.C. Y.Z., J.G. and Y.C. developed the conceptual framework for the study. C.Y., Y.Z. and W.X. provided the necessary resources. J.Y. conceived and conducted the experiments with the help of X.W., A.H. constructed the model and analysed the results. The whole process was supervised by C.Y., G.Z. Y.Z., J.G. and M.C.. J.Y., A.H. and M.C. contributed to the writing of this manuscript. All authors reviewed the manuscript.

